# The Kaposi’s sarcoma-associated herpesvirus TBP mimic uses a non-canonical DNA binding mode to promote viral late gene transcription

**DOI:** 10.1101/2025.08.25.672222

**Authors:** Lidia E. Llacsahuanga-Allcca, Allison L. Didychuk, Anthony Rodríguez-Vargas, Britt A. Glaunsinger

## Abstract

Kaposi’s sarcoma-associated herpesvirus (KSHV) orchestrates late gene transcription through viral transcriptional activators that hijack host RNA polymerase II machinery, maintaining selectivity for viral promoters. Among these, the KSHV protein ORF24 serves as a TATA-binding protein (TBP) mimic essential for recognizing viral late promoters, although the molecular mechanisms underlying its function remain poorly characterized. Here, we used AlphaFold3 to predict the structure of ORF24 in complex with DNA and validated key features in both transfected cells and during KSHV lytic replication. Structural modeling revealed that ORF24 employs a non-canonical DNA binding mode where the C-terminal domain (CTD) makes critical DNA contacts beyond the canonical TBP fold. Targeted mutagenesis confirmed that ORF24 requires conserved TBP-like phenylalanines alongside a polar-rich binding interface distinct from cellular TBP. During infection, both the TBP-like domain and CTD are essential for ORF24 occupancy at viral late promoters. Most surprisingly, we discovered that ORF24 pre-assembles with RNA polymerase II and the viral protein ORF34 to achieve stable promoter binding. This cooperative assembly mechanism represents a fundamental departure from stepwise eukaryotic transcription initiation, resembling a prokaryotic strategy within the eukaryotic nucleus.

**Summary Bullet points:**

- The structure of the KSHV TBP mimic ORF24 binding DNA was modeled and experimentally tested.
- KSHV ORF24 uses an extended DNA-binding interface beyond the canonical TBP fold.
- ORF24 requires cooperative pre-assembly with transcriptional machinery before DNA engagement

## Introduction

The gammaherpesvirus Kaposi’s sarcoma-associated herpesvirus (KSHV) is a double-stranded DNA virus that causes Kaposi’s sarcoma and other lymphoproliferative disorders, particularly in immunocompromised individuals (1,2). Like many DNA viruses, KSHV orchestrates a temporally regulated gene expression cascade during lytic infection by hijacking host transcriptional machinery. This cascade begins with Immediate Early (IE) genes followed by Early (E) genes that enable viral genome replication and culminates with Late (L) genes, which are expressed only after viral DNA replication has commenced and generally encode proteins critical for virion assembly and maturation. IE and E genes contain conventional promoter elements, suggesting their transcription proceeds in a host-like manner through the recruitment of cellular TATA-binding protein (TBP), general transcription factors (GTFs), and RNA Polymerase II (RNAPII). In contrast, the minimalistic late gene promoters, consisting of approximately 12-15 base pairs (bp), lack recognizable promoter elements beyond a distinctive TATTWAA motif that diverges from the canonical TATA box recognized by cellular TBP (3,4).

This difference in promoter architecture reflects a fundamental divergence in transcriptional mechanisms. A core eukaryotic transcription factor is TBP, which conventionally binds the TATA box sequence through a predominantly hydrophobic interface that contacts the DNA minor groove. Four conserved phenylalanine residues in TBP intercalate between DNA base pairs and induce a characteristic ∼80° DNA bend (5,6). As a core component of TFIID, TBP recruits other factors, including TFIIB, which directly bridges TBP and RNAPII to form the pre-initiation complex. Late gene expression in the beta- and gammaherpesviruses instead employs an alternative system of virus-encoded transcriptional activators (vTAs) (7,8). KSHV encodes six vTAs – ORF18, ORF24, ORF30, ORF31, ORF34, and ORF66 – that form a complex essential for late gene transcription. Among these, ORF24 plays a central role as a viral mimic of cellular TBP, recognizing the TATT motif and functionally replacing TBP on late promoters (9,10).

Previous studies have identified key protein-protein interactions within the vTA complex (11–14). ORF24 directly binds RNAPII through a conserved leucine stretch in its N-terminal domain (NTD) that interacts with hexapeptide repeats in the Rpb1 subunit (9,15). ORF24 also directly interacts with ORF34, which serves as a scaffold connecting to other vTAs (ORF18, ORF31, and ORF66) (13,16). Although site-directed mutagenesis has identified residues crucial for these interactions (including R328 for ORF34 binding), the structural basis of ORF24 function remains poorly understood, as low expression levels have made comprehensive biochemical characterization challenging. Consequently, fundamental questions remain about the function of other regions of ORF24, the molecular mechanisms governing its interactions with both viral and host factors, and how it accomplishes specificity for viral and not host promoters.

Here, we leveraged AlphaFold 3 to predict the structure of ORF24 and identify previously uncharacterized putative DNA-contacting residues, particularly within its C-terminal domain (CTD). Through systematic mutational analysis in both transfection systems and infected cells, we demonstrate that ORF24 possesses an expanded DNA-binding interface that extends beyond its TBP-like domain to include conserved CTD residues essential for late promoter recognition. We further reveal that ORF24 requires simultaneous interaction with both RNAPII and ORF34 to achieve stable late gene promoter binding, suggesting a mechanism where multiprotein complex assembly precedes stable DNA engagement. This assembly-first binding model represents a fundamental departure from classical stepwise eukaryotic transcription initiation and reveals how KSHV has evolved unique strategies to commandeer the host transcriptional machinery while ensuring selectivity for viral late promoters.

## Materials and methods

### Structural modeling and sequence analysis

Structural predictions of KSHV ORF24 and interacting proteins were generated using the AlphaFold3 server (https://alphafoldserver.com/) with amino acid sequences obtained from UniProt (accession numbers: F5HFD2 (ORF24), Q2HR98 (ORF34), P24928 (human Rpb1), Q00403 (human TFIIB)). A 30-nucleotide double-stranded DNA sequence derived from the KSHV K8.1 late promoter was included in modeling (5’-TCCGGCAGCAA**TATTAAA**GGGACCGAAGTT-3’, with the TATTWAA motif in bold). Multiple structural predictions were generated, including ORF24 alone, ORF24-ORF34-DNA, and ORF24-ORF34-DNA-TFIIB complexes. Primary analyses were performed using the highest-confidence ORF24-ORF34-DNA complex model.

Structural visualization and analysis were performed using UCSF ChimeraX version 1.8 (https://www.rbvi.ucsf.edu/chimerax)(17), with confidence scores displayed according to AlphaFold3 pLDDT values. DNA-protein contacts were identified as residues within 4 Å of DNA (center-to-center distance) or having a Van der Waals overlap of ≥0.4 Å using ChimeraX distance measurements. The DNA-protein interface was further confirmed and visualized using DNAproDB (https://dnaprodb.usc.edu/) by uploading the AlphaFold3 structure files. DNAproDB analysis used default parameters and displayed only interactions with DNA bases, minor groove, or major groove.

ORF24 homologs in beta- and gamma-herpesviruses (n=19) were identified using NCBI BLAST searches against representative viral genomes. Multiple sequence alignments were performed using Clustal Omega implemented in SnapGene (https://www.snapgene.com/) and visualized with Jalview (18). Conservation percentages were calculated based on amino acid identity across all aligned sequences.

### Plasmids

Point mutations in ORF24 were generated by inverse PCR site-directed mutagenesis using the pCDNA4.TO-ORF24-2x-STREP template (Addgene 129742) and Phusion High-Fidelity DNA polymerase (New England Biolabs). The desired point mutations were designed into the forward mutagenic primer. PCR products were treated with DpnI (New England Biolabs) for 1 hour at 37°C to digest the methylated template, then ligated using T4 DNA ligase and T4 polynucleotide kinase (New England Biolabs) according to manufacturer’s instructions. Ligation products were transformed into chemically competent *Escherichia coli* XL1-Blue cells and selected on ampicillin-containing LB agar plates.

For lentiviral expression constructs, wild-type and mutant ORF24 sequences (L73A/L74A/L75A, R328A, N425A/N427A, and N694A/N696A) were PCR-amplified from the corresponding pCDNA4.TO-ORF24 plasmids using primers that introduced an N-terminal HA epitope tag. Amplified products were subcloned into AgeI/EcoRI-digested pLJM1-zeo and pLVX-zeo vectors using In-Fusion HD cloning (Takara Bio). Wild-type ORF24 in the pLVX-zeo backbone was used for complementation studies to achieve expression levels comparable to the mutant constructs. All constructs were verified by complete plasmid sequencing.

Plasmids for viral transcriptional activator expression in luciferase reporter assays have been previously described (4,11,12): pCDNA4/TO-ORF18-2xStrep (Addgene 120372), pCDNA4/TO-ORF24-2xStrep (Addgene 129742), pCDNA4/TO-ORF30-2xStrep (Addgene 129743), pCDNA4/TO-ORF31-2xStrep (Addgene 129744), pCDNA4/TO-2xStrep-ORF34 (Addgene 120376) and pCDNA4/TO-ORF66-2xStrep (Addgene 130953). Luciferase reporter plasmids K8.1 Pr pGL4.16 + Ori (Addgene 131038) and ORF57 Pr pGL4.16 (Addgene 120378) were used to measure late and early promoter activity, respectively. The Renilla luciferase control plasmid pRL-TK (Promega) was kindly provided by Russell Vance. Lentiviral packaging plasmids pMD2.G (Addgene 12259), pMDLg/pRRE (Addgene 12251), and pRSV-Rev (Addgene 12253) were gifts from Didier Trono. The previously published pLJM1-zeo-ORF24-3x-FLAG plasmid (Addgene 130959) was used for stable complementing cell line establishment as described below.

### Cell culture

HEK293T, iSLK-puro, and iSLK-BAC16 cells were maintained in Dulbecco’s Modified Eagle Medium (DMEM) supplemented with 10% fetal bovine serum (FBS) at 37°C in a 5% CO_2_ atmosphere. HEK293T cells stably expressing ORF24-3xFLAG were additionally maintained with 500 µg/ml zeocin. iSLK-puro cells were cultured with 1 µg/ml puromycin and 250 µg/ml G418, while iSLK-BAC16 cells were maintained with additional 1000 µg/ml hygromycin for BAC selection. The iSLK cell line carrying the KSHV genome on the bacterial artificial chromosome BAC16 has been previously characterized (19). iSLK-BAC16 ORF24.stop cells previously described (9) complemented with pLJM1-zeo-HA-ORF24 were maintained with additional 500 µg/ml zeocin.

### Cell lines establishment and BAC mutagenesis

A HEK293T cell line stably expressing ORF24-3x-FLAG was established to enable propagation of ORF24-deficient KSHV mutants. Lentiviral particles were produced by co-transfecting HEK293T cells with pLJM1-zeo-ORF24-3x-FLAG and packaging plasmids pMD2.G, pMDLg/pRRE, and pRSV-Rev using Polyjet (SignaGen). After 48 hours, virus-containing supernatant was collected, filtered through 0.45 μm filters, and used to transduce fresh HEK293T cells. For transduction, 1×10^6^ freshly trypsinized HEK293T cells were seeded in 6-well plates and spinoculated with filtered supernatant containing 8 μg/ml polybrene for 2 hours at 500×g. Transduced cells were expanded to 10-cm plates after 24 hours and selected with 500 μg/ml zeocin (Sigma) for 1 week.

KSHV Bacterial Artificial Chromosomes (BACs) containing ORF24 mutations were generated using the recombination system in BAC16 GS1783 *Escherichia coli* as previously described (25). A dual-tagged parental BAC (HA-ORF24/FLAG-ORF34) was first constructed by introducing the FLAG tag into the ORF34 locus of the previously characterized HA-ORF24 BAC16 (9), followed by introduction of individual ORF24 point mutations (L73/L74A/L75A, R328A, N425A/N427A, and N694A/N696A). BAC DNA was purified using the NucleoBond BAC 100 kit (Macherey-Nagel) and mutations were confirmed by Sanger sequencing of the targeted regions. BAC integrity was verified by restriction digest analysis using RsrII and SbfI (New England Biolabs) and comparison to the parental BAC16 restriction pattern, as well as whole BAC sequencing (Plasmidusarus).

iSLK cell lines containing mutant KSHV BACs were established by co-culture as previously described (25). HEK293T cells stably expressing ORF24-3x-FLAG were transfected with 5-10 μg of purified BAC DNA using PolyJet transfection reagent (SignaGen) according to the manufacturer’s instructions. Twenty-four hours post-transfection, transfected HEK293T cells were trypsinized and co-cultured with iSLK-puro cells at a 1:1 ratio (1-1.5×10^6^ cells each) in 10-cm dishes. After allowing 4 hours for cell adherence, lytic reactivation was induced by treatment with 25 nM 12-O-tetradecanoylphorbol-13-acetate (TPA, Sigma) and 0.3 mM sodium butyrate (Sigma) in complete DMEM. Following viral reactivation and transfer, cells were selected with 1 μg/ml puromycin, 300 μg/ml hygromycin, and 250 μg/ml G418 to establish stable iSLK-BAC cell lines. The hygromycin B concentration was increased to 500 g/ml and 1 mg/ml until all HEK293T cells died.

### Late gene reporter assay

HEK293T cells (7.5×10^5^) were seeded in 6-well plates and transfected the following day with 1 μg total DNA using PolyJet transfection reagent (SignaGen Laboratories) according to the manufacturer’s instructions. Each transfection included 125 ng of each viral transcriptional activator plasmid (pCDNA4/TO-ORF18-2xStrep, -ORF30-2xStrep, - ORF31-2xStrep, −2xStrep-ORF34, -ORF66-2xStrep) and either wild-type or mutant ORF24-2xStrep (125 ng), or empty pCDNA4/TO vector (750 ng) as a negative activation control. Transfections also contained either K8.1 Pr pGL4.16 (late promoter) or ORF57 Pr pGL4.16 (early promoter) firefly luciferase reporter (125 ng) and 25 ng pRL-TK Renilla luciferase as an internal control for transfection efficiency.

Cells were harvested 24 hours post-transfection by washing twice with PBS and incubating in 500 µl Passive Lysis Buffer (Promega) for 15 minutes at room temperature. Lysate was clarified by centrifugation at 21,000 g for 2 minutes, and 20 µl of clarified lysate was added in triplicate to a white chimney well microplate (Greiner Bio-One). Luciferase activity was measured using the Dual-Luciferase Reporter Assay System (Promega) on a Tecan M1000 microplate reader. Firefly luciferase values were normalized to Renilla luciferase to control for transfection efficiency. Fold activation was calculated as the normalized luciferase signal relative to the corresponding empty vector control for each sample. Data represent the mean ± SD of at least three independent biological replicates.

### Western Blot and co-immunoprecipitation

For all DNA transfections, HEK293T cells were plated and transfected the following day at ∼70% confluency using PolyJet reagent (SignaGen) according to the manufacturer’s instructions. For protein isolation from KSHV-infected cells, iSLK cells were reactivated for 48 hours with 1 mM sodium butyrate and 5 µg/ml doxycycline in complete DMEM. Cells were washed twice with ice-cold PBS and collected with cell scrapers (Sigma-Aldrich) and pelleted by centrifugation at 500 g x 3 minutes. Cells were resuspended in RIPA lysis buffer complemented with protease inhibitor cocktail (Roche) and incubated for 30 minutes at 4°C with rotation. Protein concentrations were measured using Bradford Protein Assay (Thermo Scientific).

For immunoprecipitations, 1-2 mg of protein lysate was incubated overnight with pre-washed MagStrep type 3 XT beads (IBA Lifesciences) for strep-tagged proteins or anti-HA magnetic beads (Pierce) for HA-tagged proteins, in IP buffer (150 mM NaCl, 50 mM Tris, pH 7.4). Beads were washed three times in IP wash buffer (150 mM NaCl, 0.05% NP-40, 50 mM Tris, pH 7.4) and proteins eluted with 2X Laemmli Buffer (BioRad).

Samples were resolved by SDS-PAGE and transferred to PVDF membranes (BioRad). Membranes were blocked with 5% milk for 1 hour and incubated overnight at 4°C with primary antibodies diluted in 0.5% milk in TBST. The following primary antibodies were used: Strep-Horseradish peroxidase (HRP) (1:2500, Millipore), rabbit anti-HA (1:1000, Cell Signaling), rabbit anti-FLAG (1:1000, Cell Signaling), rabbit anti-vinculin (1:1000, Cell Signaling), rabbit anti-Rpb1 N-terminal domain (1:1000, Cell Signaling)) or mouse anti-Rpb1 (1:500, Santa Cruz), rabbit anti-K8.1 (1:10,000), rabbit anti-ORF59 (1:1000). Following incubation with primary antibody, membranes were washed three times with TBST and incubated with secondary antibody for 1h (except strep antibody). We used goat anti-rabbit immunoglobin-HRP (1:5000, Southern Biotech), and goat anti-mouse immunoglobin HRP (1:5000, Southern Biotech). Proteins were detected using chemiluminescence.

### Immunofluorescence

KSHV-infected iSLK cells were seeded on glass-bottom plates (Cellvis) pre-treated with poly-L-lysine and reactivated for 48 hours with 1 mM sodium butyrate and 5 μg/ml doxycycline in complete DMEM. Cells were washed twice with PBS and fixed with 4% paraformaldehyde in PBS for 15 minutes at room temperature. Following fixation, cells were washed with PBS and permeabilized with 0.1% Triton X-100 in PBS for 10 minutes. Cells were blocked with blocking buffer (3% BSA, 0.5% Tween-20, 0.1% Triton X-100 in PBS) for 30 minutes at room temperature with gentle rocking. Primary antibodies were diluted in blocking buffer and incubated for 1-2 hours at room temperature with rocking: rabbit anti-HA (1:500, Cell Signaling #3724) and mouse anti-FLAG (1:500, Sigma #F1804). Cells were washed three times with PBS (10 minutes each) and incubated for 1 hour at room temperature with secondary antibodies diluted in blocking buffer: goat anti-rabbit Alexa Fluor 546 (1:1,000, Invitrogen) and goat anti-mouse Alexa Fluor 647 (1:1,000, Invitrogen). Following three PBS washes (10 minutes each), nuclei were counterstained with Hoechst 33342 (1:2,500, Abcam) added to the final wash step. Cells were imaged using a Nikon spinning disk confocal microscope with a 60x oil immersion objective. Z-stack images were acquired, and representative single optical sections are shown. Image analysis was performed using Fiji.

### ChIP-qPCR

ORF24.stop BAC16 iSLK cells (3-3.5×10^6^) transduced with WT or mutant ORF24 were seeded on 15-cm plates and reactivated for 48 hours with 1 mM sodium butyrate and 5 μg/ml doxycycline in complete DMEM. Cells were cross-linked in 2% formaldehyde for 10 min at room temperature, quenched in 0.125 M glycine for 5 min, and washed twice with ice-cold PBS. Cross-linked cells were pelleted by centrifugation at 300 g for 3 minutes and stored at −80°C or processed immediately. Cell pellets were resuspended in 1 ml ice-cold ChIP lysis buffer (50 mM HEPES pH 7.9, 140 mM NaCl, 1 mM EDTA, 10% glycerol, 0.5% NP-40, 0.25% Triton X-100) supplemented with protease inhibitor cocktail (Roche), incubated at 4°C for 10 minutes with rotation, and centrifuged at 1,700×g for 5 minutes at 4°C. Nuclear pellets were resuspended in wash buffer (10 mM Tris-HCl pH 7.5, 100 mM NaCl, 1 mM EDTA pH 8.0, protease inhibitor cocktail) and rotated for 10 minutes at 4°C. After centrifugation at 1,700×g for 5 minutes at 4°C, nuclei were gently rinsed twice with shearing buffer (50 mM Tris-HCl pH 7.5, 10 mM EDTA, 0.1% SDS) with centrifugation between washes. Nuclei were resuspended in 1 ml shearing buffer and transferred to milliTUBE AFA Fiber tubes (Covaris). Chromatin was sheared using a Covaris S220 ultrasonicator for 5 minutes (peak power 140 W, duty cycle 5%, cycles per burst 200). Fragment sizes were verified using an Agilent TapeStation. Sheared chromatin was centrifuged at 16,000×g for 10 minutes at 4°C and the supernatant retained.

Chromatin was precleared with protein A/G magnetic beads (Thermo Fisher) pre-blocked with 200 μg/ml glycogen, 200 μg/ml BSA, and 200 μg/ml *E. coli* tRNA for 2 hours at 4°C. Precleared chromatin (10 μg) was diluted in shearing buffer to 500 μl, adjusted to 150 mM NaCl and 1% Triton X-100, and incubated overnight at 4°C with 10 μg anti-HA antibody (Cell Signaling #C29F4) or 10 μg rabbit IgG control (Southern Biotech). Immune complexes were captured by incubation with 25 μl pre-blocked protein A/G beads for 2 hours at 4°C with rotation. Beads were washed sequentially for 10 minutes each at 4°C with rotation: low-salt buffer (20 mM Tris pH 8.0, 1% Triton X-100, 2 mM EDTA, 150 mM NaCl, 0.1% SDS), high-salt buffer (20 mM Tris pH 8.0, 1% Triton X-100, 2 mM EDTA, 500 mM NaCl, 0.1% SDS), lithium chloride buffer (10 mM Tris pH 8.0, 0.25 M LiCl, 1% NP-40, 1% deoxycholic acid, 1 mM EDTA), and TE buffer (10 mM Tris-HCl pH 8.0, 1 mM EDTA). DNA was eluted using 100 μl elution buffer (150 mM NaCl, 50 μg/ml proteinase K) with incubation at 55°C for 2 hours followed by 65°C for 12 hours to reverse cross-links. DNA was purified using the Oligo Clean & Concentrator kit (Zymo Research) and quantified by qPCR using PowerUp SYBR Green Master Mix (Thermo Fisher). Primer sequences for target promoters are listed in Supplementary Table 1. Each sample was normalized to its own input.

## Results

### AlphaFold 3 predicts an extended DNA binding domain for KSHV ORF24

KSHV ORF24 is thought to bind viral late gene promoters through a TBP-like domain and is required for their transcriptional activation (9,10) (Figure 1A), although its structure has yet to be experimentally determined. Furthermore, only one mutant (N425A/N427A) has been experimentally confirmed as essential for its DNA binding activity (9). To better mechanistically define ORF24’s interaction with DNA, we employed AlphaFold 3 (20) to predict the structure of ORF24 in complex with its vTA binding partner ORF34 and a minimal DNA fragment of 30 bp corresponding to a late gene promoter sequence (Figure 1B, Supplementary Figure 1A). AlphaFold 3 generated a high confidence prediction (pLDDT>70) of the structural architecture of ORF24 comprising four distinct domains: 1) an N-terminal domain (NTD, residues 1-210), containing the Rpb1-binding amino acids L73-L75; 2) a central domain (residues 211-345) containing the experimentally validated ORF34-interaction residue R328; 3) the vTBP domain (residues 417-594); and 4) a C-terminal domain (CTD, residues 595-752). A low-confidence helical loop was also predicted around residues 346-416.

**Figure 1:**
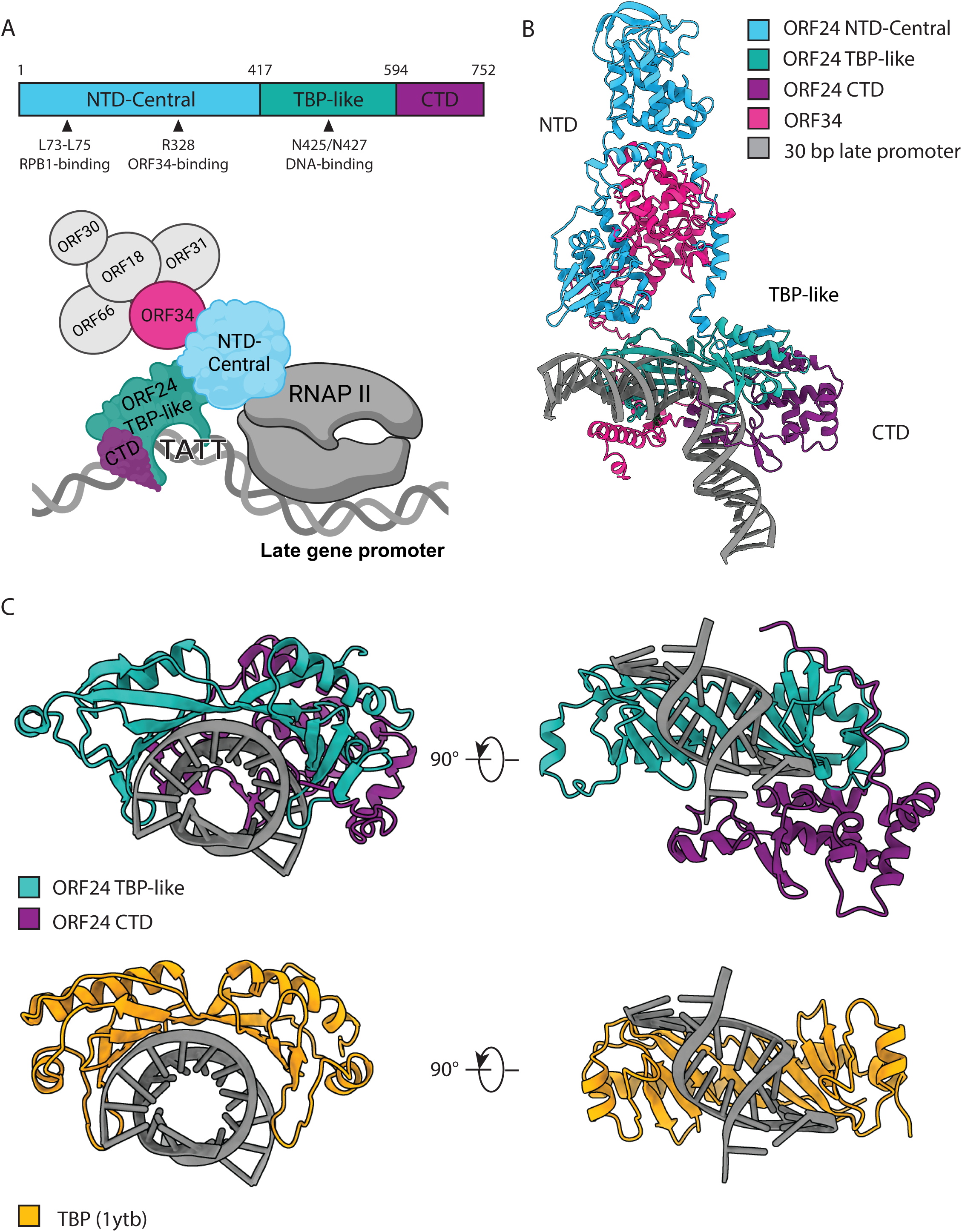
ORF24 is a viral mimic of TBP predicted to interact with DNA through conserved and novel mechanisms. A) Domains of ORF24 with relevant residues highlighted and a schematic of the vTA complex with RNA Polymerase II (RNAP II). B) AlphaFold3 predicted ORF24 structure interacting with DNA and ORF34. C) vTBP and CTD domain of ORF24 predicted to interact with DNA (top) in comparison with the known yeast TBP structure (1ytb) interacting with DNA (bottom).

AlphaFold3 predicted ORF34’s N-terminal domain (1-100 aa) with low confidence, but the rest of the protein showed moderate confidence and aligned with known interaction domains. Structural predictions including Rpb1 revealed no confident contacts with ORF24 (not shown), likely because ORF24 interacts with Rpb1 through flexible heptapeptide repeats in its disordered C-terminal tail (15), which are difficult to model computationally. These observations support the predicted organization of the complex while highlighting the challenges of modeling structurally dynamic interactions.

We then examined ORF24 conformation when bound to DNA and compared it to pre-existing knowledge of TBP and its DNA-binding mode. Intriguingly, we found that the model predicted that – in addition to the vTBP domain – the ORF24 CTD is also in close contact with the DNA backbone of the late gene promoter (Figure 1C). We used DNAproDB (21,22) to illustrate which ORF24 residues are predicted to interact with DNA in our predicted structure (Figure 2A, Table 1) and compared it to the known yeast TBP interactions (Supplementary Figure 2). This confirmed some structural similarities between the ORF24 vTBP domain and TBP, including the prediction that ORF24 forms a characteristic DNA bend through interactions with the minor groove, as well as several conserved phenylalanine, asparagine, and proline residues (Figure 1C, Supplementary Figure 2, Table 1). Notably, our analysis also confirmed previously unanticipated interactions between DNA and the ORF24 CTD (Table 1), suggesting that ORF24 has an extended DNA-binding interface that could contribute to its stable association with the late gene promoter. Consistent with this prediction, sequence alignment of ORF24 homologs across β- and γ-herpesviruses revealed that the CTD is highly conserved, particularly at several polar and positively charged residues that may support interactions with DNA (Supplementary Figure 3, Table 1).

**Figure 2:**
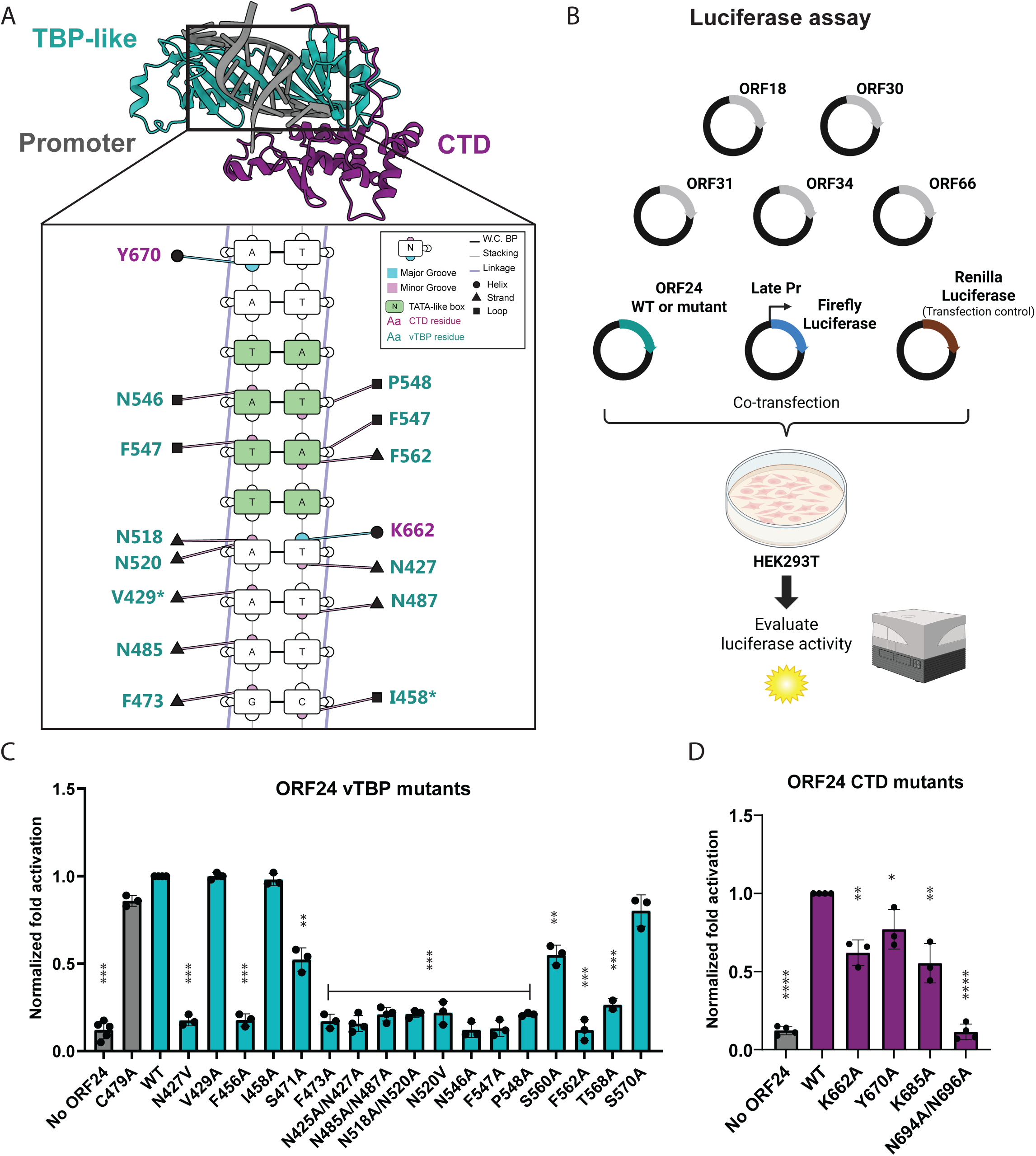
ORF24 residues predicted to bind DNA are necessary for late gene expression. A) ORF24 residues predicted to interact with the late promoter, visualized by DNAproDB. The conserved TATT motif is highlighted in green, and the residues without a significant late gene defect are marked with an asterisk (*). B) Luciferase assay schematic used to test ORF24 activity in transfected cells. C) Normalized late gene activation of ORF24 vTBP mutants after transfection in HEK293T cells. C479A ORF24 mutant was used as a negative control. D) Normalized late gene activation of CTD mutants. Data are from three independent biological replicates, with statistics being calculated using an unpaired t test. *** P 0.001; ** P 0.01; * P 0.05.

**Table 1:** ORF24 residues predicted to be important for DNA-binding.

| N | ORF24 domain | ORF24 Residue | Corresponding TBP residue | Conservation in β- and γ-Herpesviruses | Source |
| --- | --- | --- | --- | --- | --- |
| 1 | vTBP | N427 | N69 | 100% | TBP, (10) |
| 2 | vTBP | V429 | V71 | 100% | TBP, structure prediction |
| 3 | vTBP | F456 | F99 | 100% | TBP, (10) |
| 4 | vTBP | I458 | A101 | 62% | TBP, structure prediction |
| 5 | vTBP | S471 | L114 | 100% | TBP, (10) |
| 6 | vTBP | F473 | F116 | 100% | TBP, structure prediction, (10) |
| 7 | vTBP | N485 | V122 | 100% | TBP, structure prediction |
| 8 | vTBP | N487 | T124 | 100% | TBP, structure prediction |
| 9 | vTBP | N518 | N159 | 100% | TBP, structure prediction, (10) |
| 10 | vTBP | N520 | V161 | 100% | TBP, structure prediction, (10) |
| 11 | vTBP | F547 | F190 | 100% | TBP, structure prediction, (10) |
| 12 | vTBP | P548 | P191 | 100% | TBP, structure prediction, (10) |
| 13 | vTBP | S560 | L205 | 69% (S in $\gamma$, G in $\beta$) | TBP |
| 14 | vTBP | F562 | F207 | 100% | TBP, structure prediction, (10) |
| 15 | vTBP | T568 | V213 | 65% (T in $\gamma$, I in $\beta$) | TBP |
| 16 | vTBP | S570 | T215 | 69% (S in $\gamma$, P in $\beta$) | TBP, (10) |
| 17 | vTBP | N546 | L189 | 100% | Structure prediction |
| 18 | CTD | K662 | NA | 92% | Structure prediction |
| 19 | CTD | Y670 | NA | 51% | Structure prediction |
| 20 | CTD | K685 | NA | 100% | Structure prediction |
| 21 | CTD | N694 | NA | 100% | Conservation |
| 22 | CTD | N696 | NA | 100% | Conservation |

### ORF24 structural prediction is incompatible with direct TFIIB binding

To more extensively compare ORF24 with cellular TBP, we analyzed conservation of key residues identified as molecular determinants of TBP function across kingdoms of life (23), using yeast TBP as the reference sequence for structural alignment (PDB: 1YTB). ORF24 retained several conserved core residues in the N-terminal lobe (4/8 residues, including both critical phenylalanines) but fewer in the C-terminal lobe (2/8 residues) (Supplementary Table 2). Notably, ORF24 lacks the conserved negatively charged residues (E184 and E188 in yeast TBP) that are critical for TFIIB interaction (23,24), which prompted us to investigate potential structural incompatibilities with TFIIB. Structural alignment of our ORF24-DNA model with the closed PIC structure (PDB: 5IYA) revealed direct steric clashes between the ORF24 CTD and TFIIB positioning. To quantify this incompatibility, we generated AlphaFold3 predictions of ORF24-ORF34-DNA complexes with and without TFIIB. Inclusion of TFIIB significantly reduced model confidence (iPTM from 0.73 to 0.46) and eliminated many predicted ORF24-DNA contacts observed in the TFIIB-free model (Supplementary Figure 1B). This suggests that, unlike TBP, ORF24 may not directly interact with TFIIB.

### ORF24 vTBP and CTD residues predicted to bind DNA are necessary for late gene activation

We next experimentally tested our structural predictions to screen for key functional residues involved in ORF24 DNA binding. Because ORF24 is poorly expressed and not very amenable to biochemical purification, we instead used an established plasmid-based luciferase reporter assay that measures vTA-induced late gene transcription (25). It involves co-transfection of the six KSHV vTAs, a firefly luciferase reporter driven by the KSHV late K8.1 promoter, and a Renilla luciferase as a transfection control into HEK293T cells (Figure 2B). While this assay does not capture the role of viral DNA replication in potentiating KSHV late gene transcription, it nonetheless effectively recapitulates the requirement for the complete, functional vTA complex to drive late gene transcription, as well as its specificity for KSHV late but not early promoters. Furthermore, this luciferase assay enables higher throughput screening of vTA mutants for functional interrogation.

We generated 17 individual ORF24 point mutants targeting residues that structural modeling predicted to interact with DNA or that are positionally conserved with known DNA-interacting residues in cellular TBP (Table 1). We also included for comparison a mutant not predicted to be at the DNA-binding interface (C479A). To ensure that observed functional defects reflect impaired DNA binding rather than protein misfolding or disrupted protein-protein interactions, we confirmed by co-immunoprecipitation that all defective mutants were expressed comparably to WT ORF24 and retained their interactions with ORF34 and the Rpb1 subunit of RNA polymerase II (Supplementary Figure 4).

Cellular TBP employs four conserved phenylalanines that intercalate into the DNA minor groove to induce the characteristic ∼80° DNA bend essential for transcriptional initiation (26). The corresponding phenylalanines in ORF24 (F456, F473, F547, F562) were absolutely required for late gene reporter activation (Figure 2C). Similarly, a conserved proline residue (P548), which in TBP contributes to sequence specificity by creating steric constraints that favor thymine at the first TATA box position (27), was also essential for ORF24 function (Figure 2C).

A striking difference between ORF24 and TBP was the polar-rich DNA-binding interface of ORF24 compared to the predominantly hydrophobic one of TBP. Six conserved asparagine residues in the ORF24 vTBP domain (N427, N485, N487, N518, N520, N546) were crucial for late gene activation, causing complete functional loss when mutated to alanine (Figure 2C). While two of these asparagines (N425, N518)correspond to the two asparagines in TBP that form hydrogen bonds with DNA bases (6,28), the remaining four correspond to hydrophobic residues in TBP (Table 1). In contrast, mutations of hydrophobic residues to alanine (I458A, V429A) caused no significant defects in late gene activation, potentially because the alanine substitution preserved the hydrophobic character of the binding surface. The mutation outside the predicted DNA binding interface (C479A) also showed no significant defect, confirming the specificity of our functional assay.

Mutations S471A and S560A resulted in moderate reductions in late gene reporter activity, while S570A showed no significant defect (Figure 2C). These serine residues correspond to hydrophobic residues in TBP that contribute to the protein’s saddle-shaped architecture (28). The intermediate phenotype suggests these polar residues contribute to ORF24 function through weaker interactions. Collectively, these results demonstrate that ORF24 function requires many residues positionally conserved in cellular TBP, as well as novel features unique to the viral protein.

We next mutated residues in the C-terminal domain of ORF24 that were either highly conserved across ORF24 homologs and/or were predicted to bind DNA in our AlphaFold 3 model (Table 1). We confirmed these mutations did not impair ORF24 protein expression or its ability to interact with ORF34 and Pol II Rbp1 (Supplementary Figure 4). Mutations of residues predicted to directly contact DNA (K662A, Y670A, K685A) resulted in intermediate reductions in late gene reporter activity (Figure 2D). In contrast, the N694A/N696A double mutant, targeting residues in a CTD loop at the interface between the vTBP domain and the CTD, completely abolished late gene expression. These results suggest that ORF24 has an extended DNA-binding domain beyond its TBP-like fold that includes both DNA-contacting residues and structurally critical residues within its CTD.

### ORF24 mutants have late gene defects in lytic infected cells

To investigate the functional significance of these ORF24 residues during the viral lytic cycle, we engineered representative mutations into the KSHV genome using the BAC16 recombineering system in iSLK renal carcinoma cells (19). We first modified a previously published BAC16 that contains an N-terminal HA tag in the endogenous ORF24 locus (4), to include a N-terminal FLAG tag in the ORF34 locus. This enabled us to simultaneously monitor endogenous ORF24 and ORF34 protein levels, interactions, and localization in their native context upon lytic reactivation. To measure the role of the DNA binding residues, we generated an ORF24 TBP-like domain N425A/N427A mutant and a CTD N694A/N696A mutant in the dual tagged KSHV BAC16. Given that ORF24 function also requires its interaction with ORF34 and Pol II Rbp1, we additionally generated ORF24 NTD point mutants previously demonstrated to disrupt each of these interactions (R328A and L73A/L74A/L75A (also known as ‘RAAAG’), respectively) (9,13). While all the ORF24 mutants were detectably expressed upon lytic reactivation of the iSLK cells, with the exception of R328A, they were present at somewhat lower levels than WT ORF24 (Figure 3A). Each mutant had a severe defect in expression of the late protein K8.1 but expressed similar levels of the early protein ORF59 (Figure 3A). Thus, conserved residues in the NTD, ORF34-interacting domain, vTBP domain and the previously uncharacterized CTD of ORF24 are critical for late gene expression during lytic KSHV replication.

**Figure 3:**
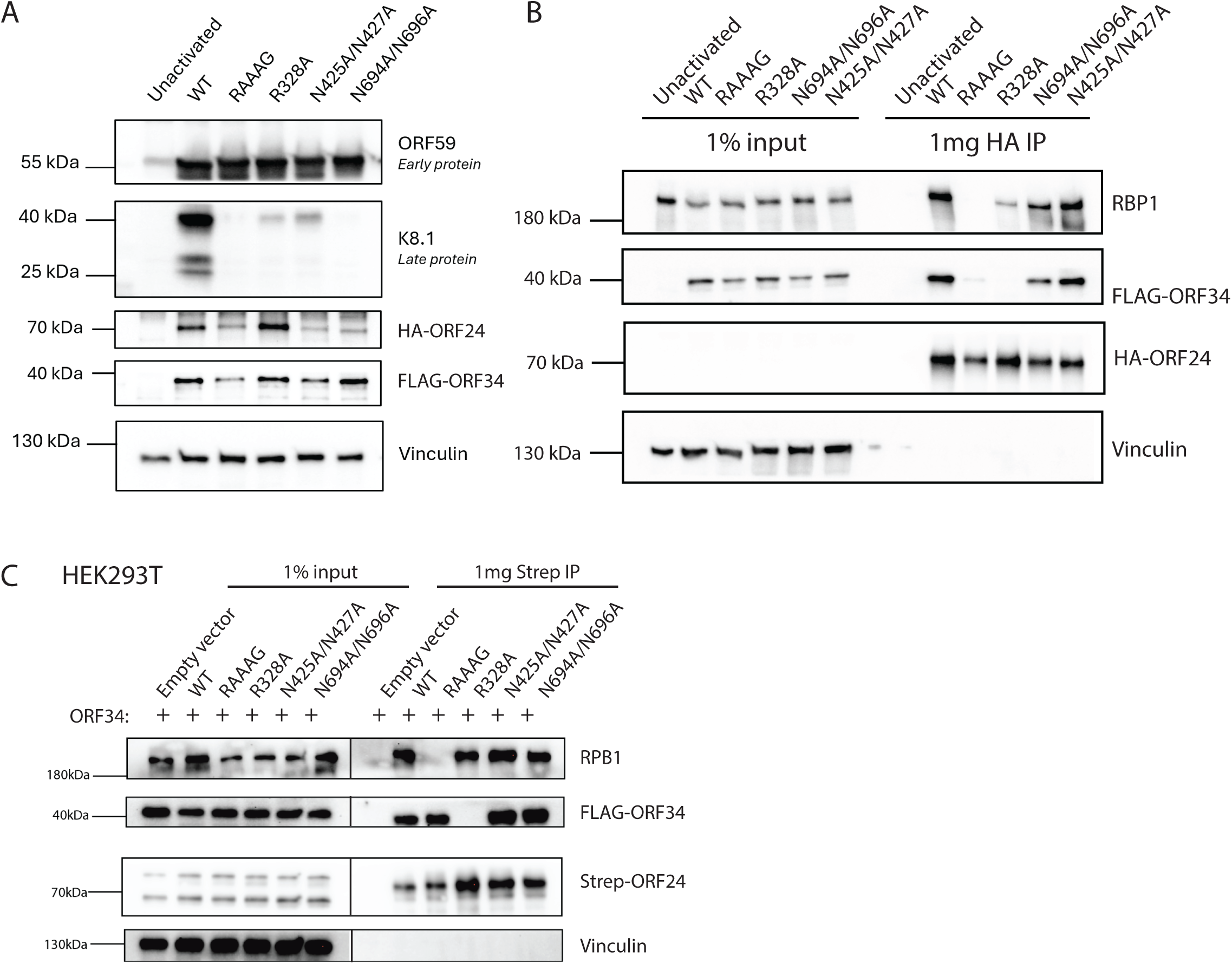
ORF24 mutants exhibit late gene defects with distinct interaction profiles during KSHV infection. A) Western blot of endogenous ORF24 mutant cell lines was performed at 48 hours post-reactivation (hpr), showing expression of a representative early (ORF59) and late (K8.1) protein, as well as protein levels of the endogenously tagged ORF24 and ORF34. Gels for ORF24 and ORF34 detection were loaded with 50 µg of protein (versus 20 µg for other proteins). B) Immunoprecipitation with HA beads was performed to enrich ORF24 in the endogenous mutant cell lines at 48 hpr. Representative Western Blot shows interaction of the ORF24 mutants with its known partners Rpb1 or ORF34. C) ORF24 mutants were transfected into HEK293T cells along with ORF34 and co-immunoprecipitated to determine its interaction partners outside of the infected context.

### ORF34 and Rpb1 co-stabilize the vPIC complex during lytic infection

We next evaluated how each of these mutations in KSHV impacted the ability of ORF24 to interact with the Rpb1 subunit of Pol II as well as with ORF34. Immunoprecipitation of HA-ORF24 at 48h post lytic reactivation confirmed that the vTBP (N425A/N427A) and CTD (N694A/N696A) DNA binding mutants of ORF24 retain the ability to bind Rpb1 and ORF34 (Figure 3B). Interestingly, however, the ORF24 Rpb1 interaction-deficient mutant (RAAAG) had a weaker interaction with ORF34 compared to WT ORF24. Correspondingly, the ORF34-interaction-deficient mutant (R328A) interacted poorly with Rpb1. This suggests that in the context of lytic viral replication, ORF24’s interactions with ORF34 and Rpb1 are both important for formation of a stable vPIC complex, such that disruption of one interaction destabilizes the other. Notably, these interaction dynamics were specific to the viral infection context and were not observed in transient 293T cell transfection experiments here (Figure 3C) and in prior publications (13).

### ORF24 mutants localize to replication compartments during lytic infection

To activate late genes, ORF24 must localize to viral replication compartments, which serve as critical hubs for KSHV DNA replication, transcription, and packaging. We next considered whether the defects in late gene transcription for any of the ORF24 mutants could be explained by a failure to localize to replication compartments. Neither ORF24 nor ORF34 have recognizable nuclear localization signals, so one hypothesis is that they are brought into the nucleus through the interaction of ORF24 with Rpb1. However, all ORF24 mutants reproducibly localized to viral replication compartments in reactivated iSLK cells when visualized by immunofluorescence staining for the HA epitope (Figure 4). FLAG-ORF34 similarly localized to nuclear replication compartments but was also found in condensed cytoplasmic puncta in late stage reactivated cells. The fact that the localization of neither ORF24 nor ORF34 is disrupted by any of the mutations indicates that localization of these proteins to nuclear replication compartments is not dependent on their interactions with each other, with DNA, or with Rpb1.

**Figure 4:**
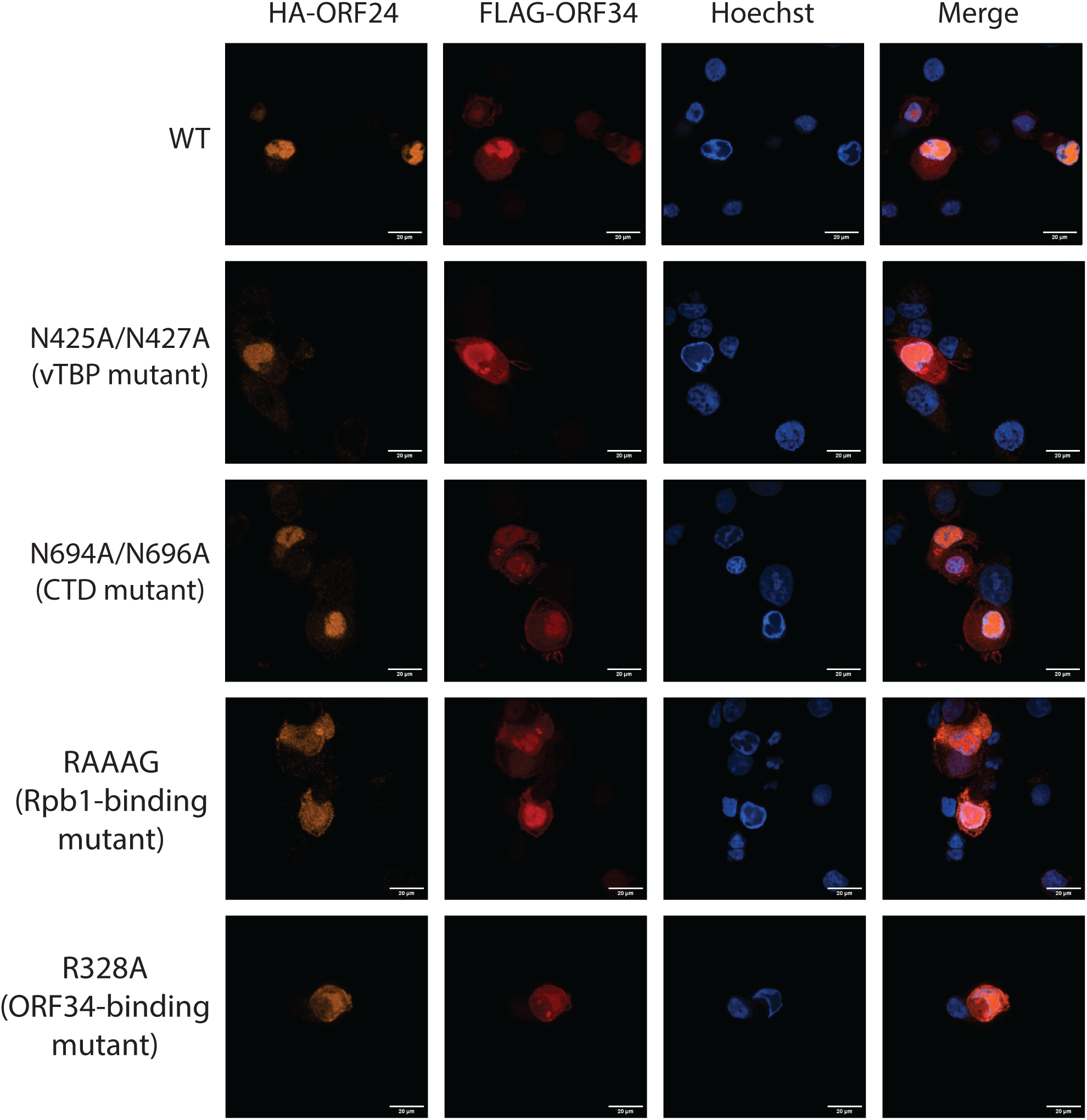
Endogenous ORF24 mutants localize to replication compartments at late times of lytic infection. iSLK cells expressing endogenous HA-tagged ORF24 mutants and FLAG-tagged ORF34 were reactivated and processed for immunofluorescence at 48 hours post-reactivation. Cell nuclei were stained with Hoechst. In late-stage reactivated cells, viral replication compartments appear as Hoechst-excluding regions within the nucleus.

### ORF24 promoter occupancy in infected cells requires multiprotein complex pre-assembly

Finally, we sought to directly measure the promoter binding capabilities of the ORF24 mutants during infection using chromatin immunoprecipitation (ChIP). However, the lower expression of the RAAG, vTBP, and CTD mutants in our endogenously tagged cell lines precluded quantitative measurements of their DNA binding (Figure 3A). To circumvent this limitation, we instead used a complementation strategy in which we stably transduced an ORF24.stop iSLK cell line with wild-type and mutant HA-ORF24 constructs. The complemented cell lines demonstrated significantly improved HA-ORF24 expression (Figure 5A). We then evaluated the ability of the complemented HA-ORF24 to bind late or early promoters using ChIP-qPCR. Wild-type HA-ORF24 selectively bound the late K8.1 promoter but not the early ORF57 promoter, demonstrating specificity for late-stage transcriptional regulation. In contrast, neither the vTBP (N425A/N427A) nor the CTD (N694A/N696A) mutant detectably bound the late gene promoter (Figure 5B), in agreement with its predicted role in DNA binding. Interestingly, both the Rpb1-interaction-deficient and the ORF34-interaction-deficient mutants also failed to bind late promoters (Figure 5B). Thus, in addition to requiring its extended DNA binding domain, ORF24 must also be in a complex with ORF34 and RNAP II to stably interact with the K8.1 late promoter. This further differentiates KSHV late gene transcription from the classical stepwise assembly model of mammalian transcription, where several general transcription factors bind the promoter prior to recruitment of RNAP II (Figure 5C).

**Figure 5:**
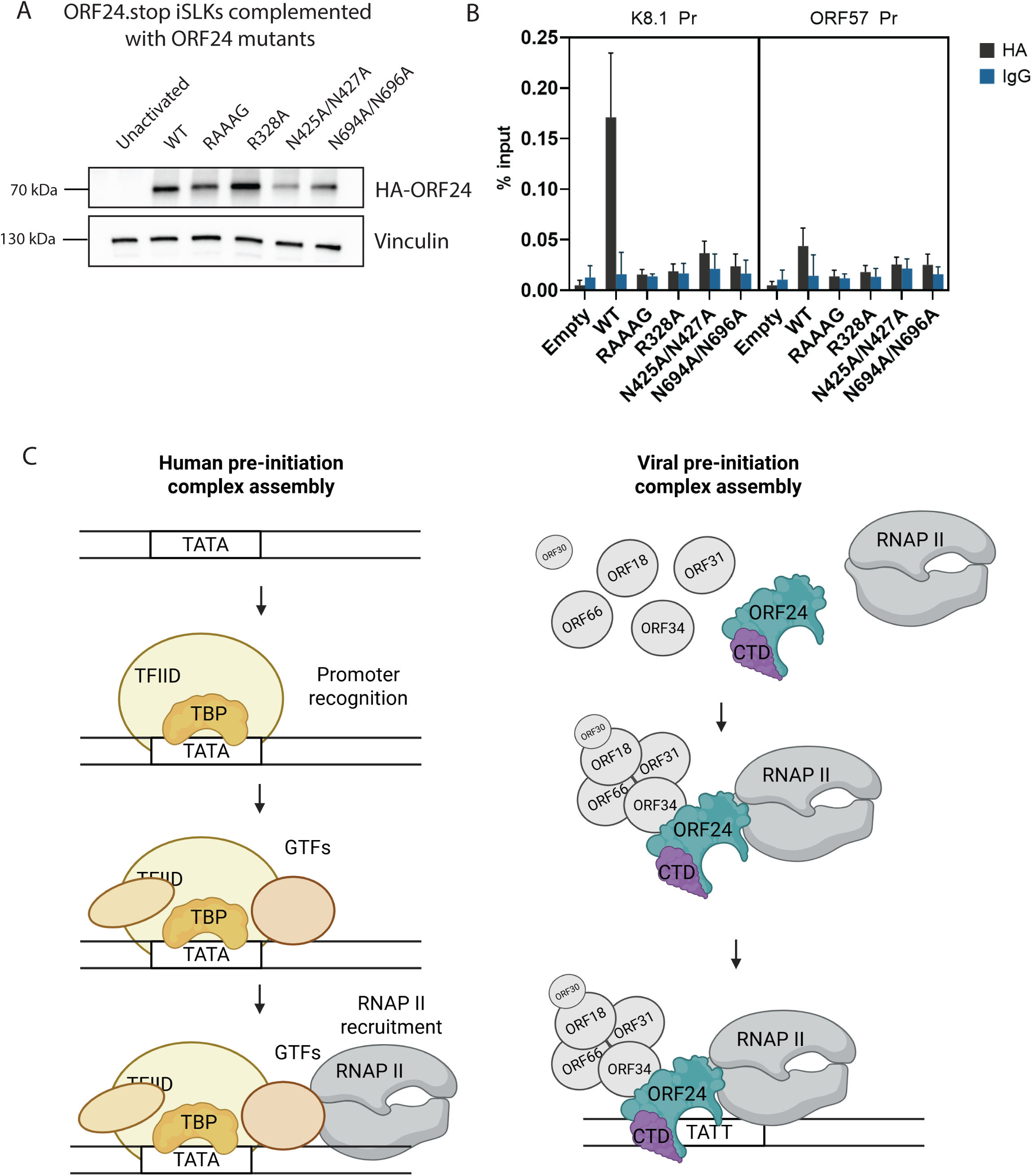
ORF24 requires interaction with Rpb1 and ORF34, as well as its CTD and vTBP to bind late promoters in infected cells. A) ORF24 protein levels at 48 hpr in complemented iSLK cell lines. B) ChIP-qPCR of indicated cell lines at 48 hpr was performed with HA-antibody, and IgG as a negative control. qPCR to detect the associated DNA was performed with promoter-specific primers for a representative late (K8.1) and early (ORF57) promoter. C) Proposed model of ORF24 recognition of late promoters in comparison to eukaryotic mechanism.

## Discussion

By integrating structural predictions from AlphaFold with experimental assays, we demonstrate how KSHV ORF24 uses a distinctive strategy for viral transcriptional control, fundamentally different from canonical eukaryotic transcription initiation pathways. Our systematic mutagenesis approach of the TBP-like domain reveals both conserved and divergent features that distinguish this viral TBP mimic from its cellular counterpart. ORF24 activity is dependent on phenylalanine residues previously described as critical molecular determinants of TBP function across all domains of life (23), suggesting conservation of the fundamental DNA-bending mechanism present in eukaryotic and archaeal TBP. Yet, the ORF24 DNA-binding interface is more polar compared to the predominantly hydrophobic interface of TBP. ORF24 function also requires six asparagine residues (compared to TBP’s dependency on only two), which commonly form hydrogen bonds with specific bases of DNA (29). This polar-rich interface likely reflects adaptation to the specific sequence and structural features of KSHV TATTWAA-containing promoters, potentially providing enhanced binding specificity or stability to viral promoters.

A distinctive feature of ORF24 is its extended DNA binding interface mediated by its CTD. Our structural predictions suggest that the CTD engages DNA through contacts distinct from the canonical minor groove interactions of the TBP-like domain. Our reporter assay confirmed intermediate late gene activation defects when we mutate the residues predicted to directly contact DNA (K662A, K685A, Y670A) and a complete loss of late gene activation in the N694A/N696A mutant, whose residues are not predicted to be in direct contact with DNA. These results suggest that the contribution of the CTD to DNA binding may be more architectural than mediated through specific key contacts with DNA: The N694/N696 residues may serve as critical structural elements that maintain the proper conformation of the CTD for DNA engagement, and the predicted DNA-contacting residues may provide stabilizing interactions. The functional importance of the CTD is further supported by its high sequence conservation across β- and γ-herpesviruses, particularly at positively charged residues that may facilitate DNA interactions, suggesting this extended DNA binding mechanism is likely conserved across these herpesvirus subfamilies. While we confirmed the requirement for the CTD in DNA binding during infection, direct biochemical validation of the predicted DNA contacts and structural studies will be necessary to explore this novel DNA binding mode.

Interestingly, ORF24 lacks the conserved negatively charged residues critical for TFIIB interaction and our structural modeling suggests spatial conflicts with TFIIB positioning. This indicates that, unlike TBP, ORF24 might not directly interact with TFIIB. Consistent with our observations, computational modeling of other β- and γ-herpesvirus transcriptional complexes has suggested that other vTAs like ORF31 and its homologs may also be incompatible with canonical TFIIB positioning (30). While TFIIB has been detected at late promoters (9), this may not reflect simultaneous binding with ORF24 at individual promoter molecules. Whether these factors engage promoters in a temporally distinct manner or whether TFIIB plays a different role in viral transcription remains to be determined.

In eukaryotic transcription, TBP can bind promiscuously (31) and other GTFs provide specificity and regulation (32,33). The unique ORF24 characteristics of a polar-rich DNA binding interface and extended DNA binding domain may be an evolutionary adaptation to provide specificity to viral promoters within a simplified viral PIC. This architecture together with other vTAs may functionally replace multiple cellular GTFs in viral late transcription. Determining which specific GTFs are supplanted by viral factors represents an important avenue for future investigation.

Analysis of ORF24 mutants in KSHV-infected cells revealed a synergistic interaction between members of the viral PIC, where impaired binding to either RNAP II or ORF34 resulted in proportionally greater loss of overall complex stability than would be expected from individual defects alone. This could occur if the vTA complex exists as an interdependent assembly where interaction with either partner stabilizes interaction with the other through conformational changes. Alternatively, incomplete complexes may be subject to enhanced degradation. Cellular quality control mechanisms are known to target misfolded or incompletely assembled protein complexes for proteasomal degradation (34), which could explain the lower expression of mutant ORF24 proteins.

A key finding of our study is that ORF24 requires intact interactions with both RNAP II and ORF34 for stable promoter binding, demonstrating an assembly-first mechanism where multiprotein complex formation precedes DNA engagement. Previous work had shown the requirement of other vTAs (ORF66 and ORF30) for ORF24 DNA engagement (11), highlighting the importance of an intact vTA complex. Together with our ORF34 binding-defective mutant, these data suggest a requirement for the complete vPIC before DNA engagement. Surprisingly, our work demonstrates that RNAP II is also pre-assembled before DNA binding. This represents a fundamental departure from the canonical stepwise model of eukaryotic transcription initiation and may help prevent inappropriate binding to cellular TATA-containing sequences. The requirement for pre-initiation complex formation effectively restricts ORF24 activity to viral transcriptional contexts, providing an additional layer of selectivity beyond sequence recognition alone.

Our work establishes ORF24 as a sophisticated viral adaptation that consolidates multiple transcriptional functions—TBP-like DNA recognition, extended DNA binding, and assembly-dependent specificity—into a single polypeptide. This streamlined system resembles bacterial sigma factors more than eukaryotic transcription complexes, reflecting how viruses have evolved elegant solutions to commandeer host transcriptional machinery while maintaining specificity. The assembly-first mechanism we describe for ORF24 provides a tractable model for understanding cooperative binding principles that may be obscured by redundancy in cellular systems. As single-molecule studies continue to reveal the dynamic nature of transcriptional regulation, viruses like KSHV offer powerful platforms for dissecting fundamental mechanisms of transcriptional control.

## Data availability

The data underlying this article are available in the article and in its online supplementary material.

## Funding

This research was supported by NIH grant R01AI122528 to B.G., who is also an investigator of the Howard Hughes Medical Institute.

## Acknowledgements

We would like to thank all members of the B.G. lab for helpful discussions and suggestions, as well as reading of the manuscript. Confocal imaging experiments were conducted at the CRL Molecular Imaging Center, RRID:SCR_017852, supported by the Gordon and Betty Moore Foundation.

## Notes

### Competing Interest Statement

The authors have declared no competing interest.

### Summary of Updates

Results and discussion sections updated to mention the conservation of the CTD of ORF24. Supplementary data updated to include corresponding analysis. Figure 2 revised.

