## Supplemental Data for "The Kaposi’s sarcoma-associated herpesvirus TBP mimic uses a non-canonical DNA binding mode to promote viral late gene transcription"

### Supplementary Data

**Supplementary Table 1:** Primers used for ChIP-qPCR

| **Target** | **Forward Primer** | **Reverse Primer** |
|---|---|---|
| K8.1 Pr | GGGAGAACCATGCCAGACTTTG | GCATAGGATTAGGAGCGCCAC |
| ORF57 Pr | CAGTGTTTTGCCAGCAAGTG | GGGCTATTTTGGGAACCTG |

**Supplementary Table 2:** Alignment of ORF24 to yeast TBP (1YTB) key residues from (20), with common residues bolded

| **N** | **Category** | **Lobe** | **Common TBP Numbering (CTN)** | **Yeast_res** | **Yeast_res_nr** | **ORF24_res** | **ORF24_res_nr** |
|---|---|---|---|---|---|---|---|
| 1 | core | N | **L3.27** | **F** | **99** | **F** | **456** |
| 2 | core | N | S4.5 | I | 115 | V | 472 |
| 3 | core | N | **L5.1** | **F** | **116** | **F** | **473** |
| 4 | core | N | **L5.3** | **S** | **118** | **S** | **475** |
| 5 | core | N | **L5.4** | **G** | **119** | **G** | **481** |
| 6 | core | N | S5.1 | K | 120 | T | 482 |
| 7 | core | N | S5.6 | G | 125 | N | 487 |
| 8 | core | N | L6.1 | K | 127 | S | 489 |
| 9 | core | C | **L3.27** | **F** | **190** | **F** | **547** |
| 10 | core | C | S4.5 | I | 206 | W | 561 |
| 11 | core | C | **L5.1** | **F** | **207** | **F** | **562** |
| 12 | core | C | L5.3 | S | 209 | A | 564 |
| 13 | core | C | L5.4 | G | 210 | A | 565 |
| 14 | core | C | S5.1 | K | 211 | T | 566 |
| 15 | core | C | S5.6 | G | 216 | K | 571 |
| 16 | core | C | L6.1 | K | 218 | Y | 573 |
| 17 | TFIIB-binding | C | L3.2 | E | 186 | W | 543 |
| 18 | TFIIB-binding | C | L3.4 | E | 188 | T | 545 |
| 19 | Regulatory | N | L2.3 | R | 90 | P | 450 |
| 20 | Regulatory | N | L2.4 | N | 91 | R | 451 |
| 21 | Regulatory | N | H2.13 | R | 137 | F | 495 |
| 22 | Regulatory | N | H2.17 | R | 141 | M | 499 |
| 23 | Regulatory | N | H2.21 | K | 145 | K | 503 |

Supplementary Figure 1: Confidence scores of predicted models A) Top model of ORF24, ORF34 and DNA predicted by AlphaFold3 colored by its confidence score. B) AlphaFold3 predicted model of ORF24-ORF34-TFIIB and 30 bp of a late promoter (left), the TBP-like domain of ORF24 highlighted in teal in this predicted structure (right).

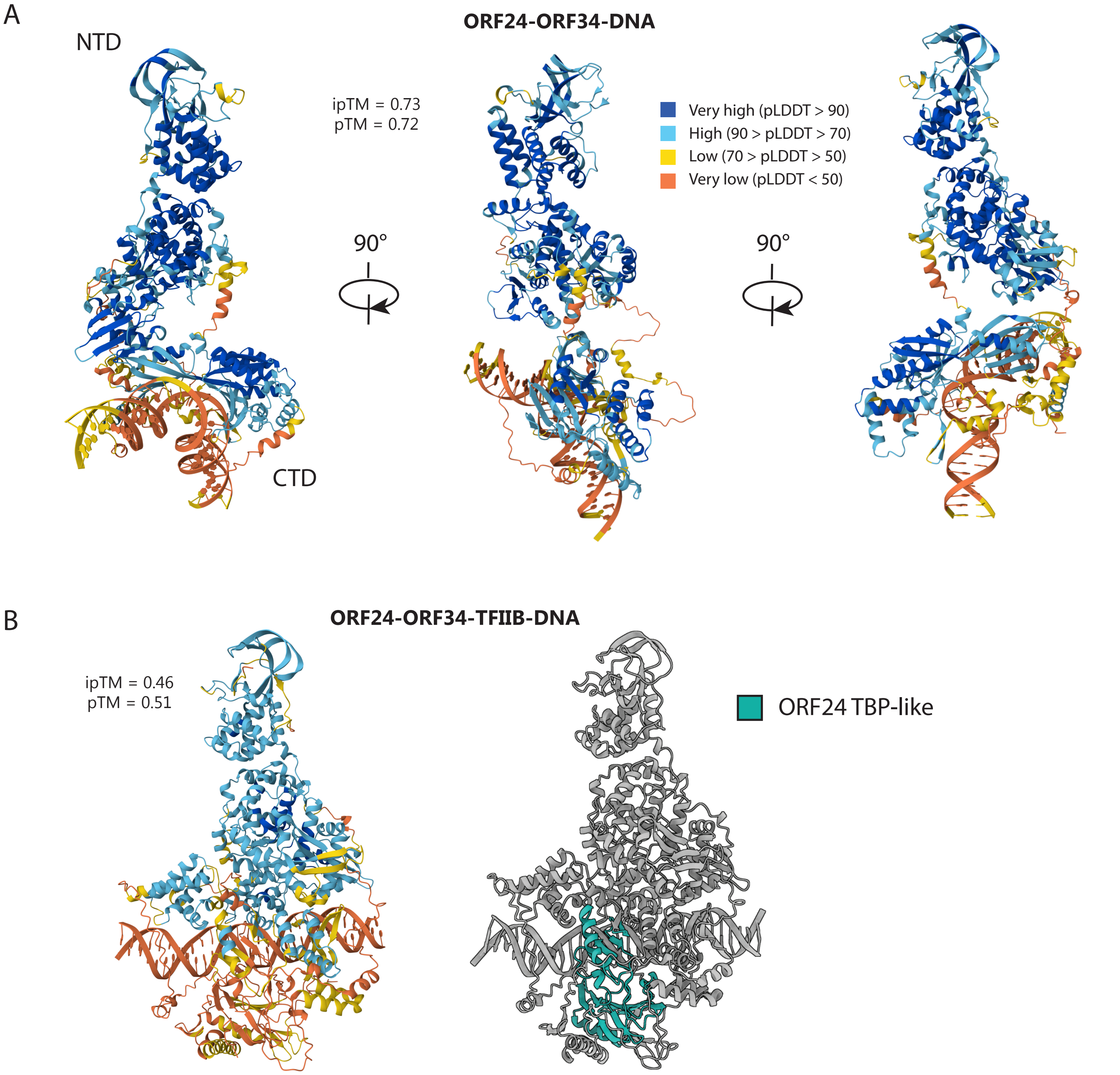

Supplementary Figure 2: Comparison of protein-DNA interactions of yeast TBP (PDB: 1ytb) and our predicted ORF24-ORF34-DNA model. The interactions showed here are predicted and visualized by DNAproDB. Conserved Phenylalanines are colored in orange, Asparagines are shown in blue, and the conserved Proline is shown in red. ORF24's unique CTD residues are shown in purple.

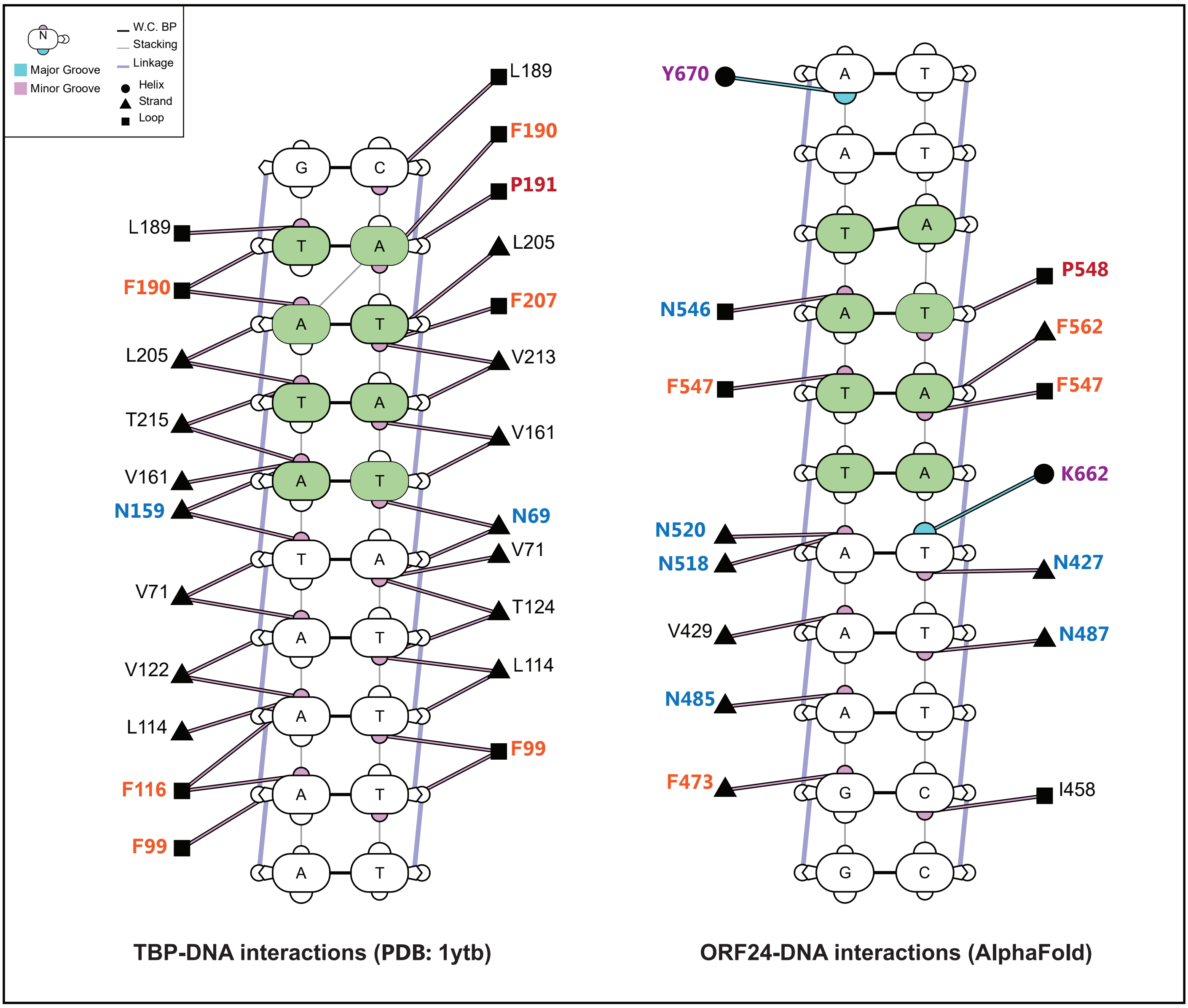

Supplementary Figure 3: Protein sequence alignment of 19 ORF24 homologs showing the C-terminal domain. Residues are colored by physical properties and conservation using Clustal, residues with at least 30% identity are colored. ORF24 is the reference sequence and the numbering on top of the sequences corresponds to it. For brevity, the alignment is truncated to show only residues corresponding to ORF24 positions 596-731, which includes the most conserved residues of the CTD.

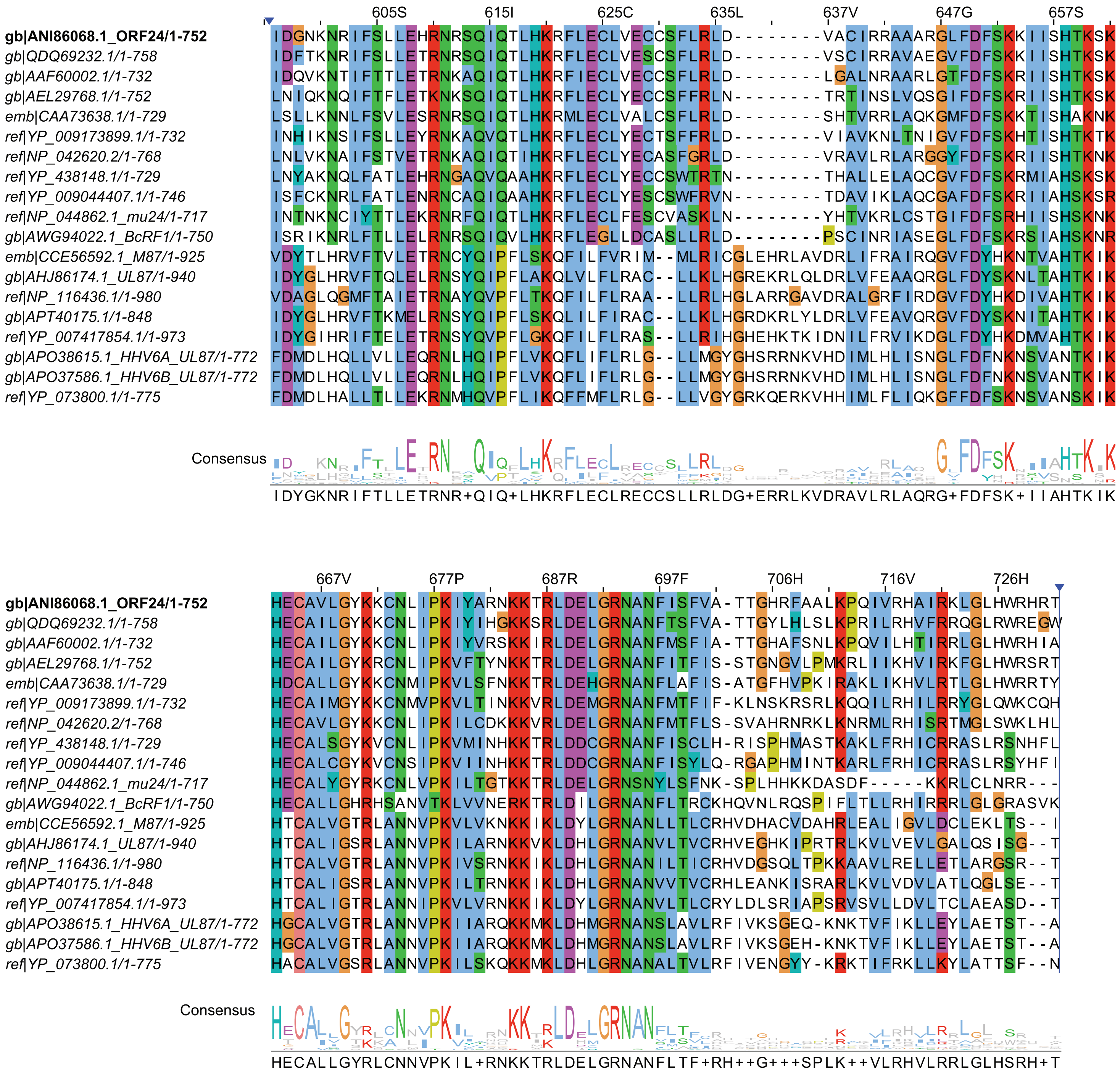

Supplementary Figure 4: Observed late gene defect in tested ORF24 mutants is not due to lower expression or defective interaction with Rpb1 and or ORF34. Co-immunoprecipitation after transfection shows normal interactions and expression of all tested ORF24 mutants. The star (*) in the input highlights unspecific bands of the Strep antibody. A) Asparagine double mutants from vTBP and CTD. B) Serine, Proline and Threonine mutants from the vTBP domain. C) Phenylalanine mutants. D) Evaluated CTD mutants and N546A from vTBP.

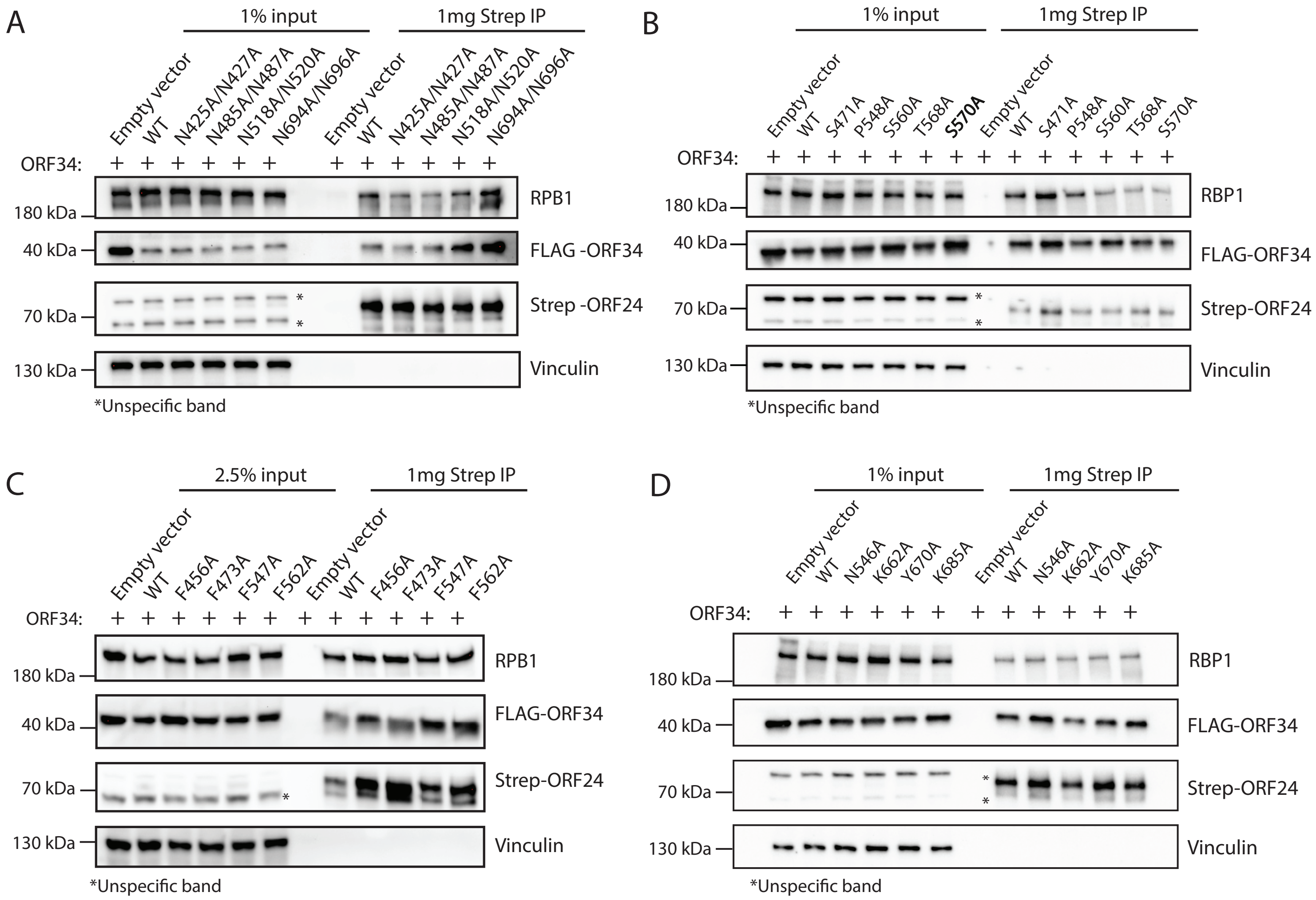
